# Reductive carboxylation via isocitrate dehydrogenase 1 supports cardiac metabolic adaptation during oncometabolic stress

**DOI:** 10.64898/2026.06.06.727699

**Authors:** Tanvi Shankar, Yaqi Gao, Zipporah Erebholo, Nicole Nakama, Kyoungmin Kim, Ian K. Williamson, Nathaniel Snyder, Evan P. Kransdorf, Ralph DeBerardinis, Heinrich Taegtmeyer, Brandon Faubert, Anja Karlstaedt

**Author notes:** **CORRESPONDING AUTHOR** Anja Karlstaedt, MD, PhD, Assistant Professor, Department of Cardiology, Smidt Heart Institute, Cedars Sinai Medical Center, 127 San Vicente Blvd, Advanced Health Science Pavilion 9229 Los Angeles, California 90048, USA.

## Abstract

**Background:** Cardiovascular disease and cancer are the two leading causes of morbidity and mortality worldwide. Metabolic dysregulation of cancer cells extends beyond the tumor microenvironment and increases the risk for cardiovascular diseases. One common somatic mutation in cancer cells affects isocitrate dehydrogenase (IDH) 1 and 2, which catalyzes the oxidative decarboxylation of isocitrate to alpha-ketoglutarate in the cytosol and mitochondria, respectively. IDH1 and 2 mutations cause the production of the oncometabolite D-2-hydroxyglutarate (D2-HG), which allosterically inhibits α-ketoglutarate dehydrogenase (α-KGDH) and is associated with reduced cardiac contractile function.

**Methods:** We combined stable isotope tracer studies with computational modeling to investigate the fundamental role of IDH isoforms in cardiac adaptation under oncometabolic stress.

**Results:** We uncovered an unexpected cardiac phenotype that expands the role of IDH1 in the heart beyond oxidative metabolism. We quantified the stable isotopomer distributions from glucose and glutamine in perfused working rat hearts and isolated adult ventricular cardiomyocytes using mass spectrometry-based metabolomics. Our analysis revealed that defective mitochondrial metabolism causes the redirection of carbon flux from oxidative towards reductive pathways. Reductive carboxylation of α-KGDH increases glutamine uptake and glutamine-derived citrate formation in working rat heart perfusions and cultured adult mouse ventricular cardiomyocytes. To identify which IDH isoform is responsible for redirecting carbon flux, we developed knockout models of IDH1, IDH2, and IDH3 in adult mouse ventricular cardiomyocytes. Loss of IDH1 expression impaired the reductive formation of citrate and caused functional defects in cardiomyocytes. Lastly, epigenetic analyses of histone marks revealed that IDH1 induces widespread alterations in histone acetylation and tri-methylation.

**Conclusion:** Our results highlight a novel role for IDH1 in cardiac metabolism and transcriptional control of metabolic adaptation to tumor-mediated stress and provide evidence that reductive-citrate formation may induce epigenetic modifications in the heart.

## INTRODUCTION

Cardiovascular disease (CVD) and cancer are the leading causes of morbidity and mortality in the U.S.^1^ Heart failure (HF) is a significant adverse cardiac outcome affecting 10 to 20% of cancer survivors and patients.^2^ The risk for cardiovascular mortality is approximately seven times higher in this patient cohort compared with the age- and sex-adjusted general population. One of the crucial factors of HF is metabolic adaptation, which results in the activation of protein and gene programs, structural cardiac remodeling, and subsequent left ventricular dysfunction. Increasing evidence has demonstrated that cancer-related inflammation^3–5^, oxidative stress^6–8^, impaired mitochondrial function^7,9^ and modified systemic metabolism are involved in the pathogenesis of cardiomyopathy^5,9–11^. Despite decades-long efforts, the mortality and incidence of cardiovascular diseases remain high and are a major cause of death among cancer patients and survivors^12^. Thus, there is an urgent need for new therapeutic targets to treat CVDs and protect the heart during cancer treatments.

The development of HF and cardiomyopathy is tightly linked with cardiac energy metabolism, and numerous studies have demonstrated that disrupting cardiac energy homeostasis leads to organ dysfunction^13–17^. Likewise, cancer cells alter their metabolism to ensure growth and proliferation^18–22^. During the progression of cancer, somatic mutations are acquired that frequently impact enzymatic function, which have been linked to declining cardiac function^9,11^. Isocitrate dehydrogenase (IDH) is a critical enzyme in the Krebs cycle that catalyzes the decarboxylation of isocitrate to α-ketoglutarate. Three isoforms have been identified, which are in the cytosol (IDH1) and mitochondria (IDH2 and IDH3). While IDH3 catalyzes the irreversible NADH-dependent step, IDH1 and IDH2 carry reversible NADPH-dependent conversions. Recurrent somatic mutations in IDH1 and IDH2 occur in approximately 80% of grade II-III gliomas and secondary glioblastoma, 10-20% of acute myeloid leukemia (AML) patients, and 5-10% of cholangiocarcinoma. Most point mutations are found in active site arginine residues of IDH1 (R132) and IDH2 (R140 and R172)^23,24^, which causes a neomorphic enzymatic function that catalyzes the conversion of α-ketoglutarate (α-KG) to the enantiomer D2-hydroxyglutarate (D2-HG)^25,26^. D2-HG is a structural homolog to α-KG and competitively inhibits Fe(II)/α-KG-dependent dioxygenases^27^, and metabolic enzymes including α-KGDH^9,10^. Previous studies have demonstrated that cancer-specific production of D2-HG impairs cardiac contractile function by inhibiting α-KGDH and disrupting oxidative metabolism in the heart. Likewise, global expression of mutant IDH2 in transgenic mice induced dilated cardiomyopathy and muscular dystrophy^11^. Furthermore, clinical studies revealed that oncometabolic production of D2-HG exacerbates doxorubicin-induced cardiac toxicities^4^.

Because IDH mutants block cell differentiation and promote tumor transformation, inhibition of IDH1 and IDH2 is a therapeutic strategy. Selective IDH inhibitors have been developed and recently received FDA approval for the treatment of AML with IDH1 mutation (AG-120, ivosidenib), IDH2 mutation (AG-221, enasidenib), and for patients with grade 2 astrocytoma or oligodendroglioma with a susceptible IDH1 or IDH2 mutation (vorasidenib). Despite these advancements in IDH-targeted cancer therapies, the roles of IDH1 and IDH2 in cardiac metabolic adaptation and remodeling during cancer remain unclear. Recent studies suggest that cancer cells promote metabolic vulnerabilities in other cell types distant from the primary tumor. Discovering key metabolic features in cardiac pathophysiology during oncometabolic stress may provide new targets and strategies for the treatment of cancer patients and HF.

In this study, we examined the role of IDH isoforms in oncometabolic stress, finding that IDH1 carries reductive formation of citrate and drives metabolic adaptation in cardiomyocytes. This phenomenon was further validated in human heart tissue from healthy donors and tracer studies combined with computational modeling. IDH1 deficiency effectively prevented the reductive formation of citrate from glutamine and increased glycolytic flux. Interestingly, our mechanistic studies showed that clinically relevant IDH1 inhibitors confer cardio-protective benefits. These studies reveal that reductive carboxylation via IDH1 is a new regulator of cardiac adaptation and heart failure.

## METHODS

### Data Availability

The data and detailed methods supporting the findings in our study are described in the Supplemental Material and Major Resources table in the online supplementary files.

### Statistical Analysis

Data were tested for normal distribution using the Shapiro-Wilk and Kolmogorov-Smirnov tests. For normal distributed data, we conducted a t-test followed by multiple-comparison analysis, with a false discovery rate of less than 5% using the two-step method of Benjamini, Krieger, and Yekutieli. For non-normally distributed data, including tracer analysis data, we conducted the Mann-Whitney test, followed by multiple comparisons analysis, with a false discovery rate (FDR) of less than 1% using the two-step method of Benjamini, Krieger, and Yekutieli. Differences between groups for other data types were tested using a t-test, one-way ANOVA, or two-way ANOVA, as appropriate. Partial Least Squares Discriminant Analysis (PLS-DA) was conducted on normalized metabolomics data using the R package mdatools (v0.14.2). The following R packages were used for general data analysis: base (v4.3.2), dplyr (v1.1.4), readxl (v1.4.3), tibble (v3.2.1), tidyverse (v2.0.0); and Data Visualization: ggplot2 (v3.5.2).

## RESULTS

### Altered citrate metabolism is a common feature of oncometabolic stress in the human heart

We analyzed metabolomics in human heart tissue slices to identify the effects of oncometabolic stress on the human heart. We generated viable agarose-embedded slices of heart tissues from six to seven normal human heart donors (**Figure 1A**). Left ventricular tissue slices were embedded in agarose, and samples were labeled with [U-^13^C]glucose *ex vivo* in a medium formulated to contain a nutrient content similar to human plasma and supplemented with or without the oncometabolite D2-HG (1.0 mM)^9,28^. Heart tissue slices were labeled for 24h and then freeze-clamped for targeted metabolomics and isotopomer distribution analysis. Heart tissue slices were randomly assigned to control or experimental groups, with tissue samples from the same patient serving as controls. PLS-DA analysis revealed a clear separation in metabolic profiles between human tissue samples treated with D2-HG and untreated controls (**Figure 1B**). Metabolic abundance of citrate and hypotaurine was significantly increased compared to untreated controls (**Figure 1C** and **Supplementary Figure 1A**). Next, we used [U-^13^C]glucose to trace the metabolic fate of glucose in human hearts during oncometabolic stress. The incorporation of glucose-derived carbons into pyruvate m+3 and lactate m+3 (corresponding with a fully labeled molecule) was increased in heart tissue samples treated with D2-HG compared to untreated controls (**Figure 1D**). Likewise, citrate m+2 and the labeling ratio of citrate m+2 to pyruvate m+3 were higher in D2-HG-treated tissue slices (**Figure 1E and Supplementary Figure 1B**), indicating an increased contribution of glucose to mitochondrial respiration and oxidative metabolism. In contrast, the malate m+2 to citrate m+2 ratio (**Figure 1F**) was significantly reduced in D2-HG-treated compared to untreated tissue slices, suggesting that glucose-derived pyruvate is converted to citrate but incompletely decarboxylated to malate. We previously demonstrated that D2-HG inhibits α-ketoglutarate dehydrogenase activity^9^, thereby disrupting the Krebs cycle flux and promoting the redirection of citrate into other pathways, including export into the cytosol and conversion into oxaloacetate and acetyl-CoA via ATP-dependent citrate lyase (ACL). Together, these data revealed an increased labeling of glycolytic and Krebs cycle intermediates from [U-^13^C]glucose during oncometabolic stress, which is consistent with our previous reports in the isolated working rat heart^9^ and supports an oncometabolic profile in the heart.

**Figure 1.**
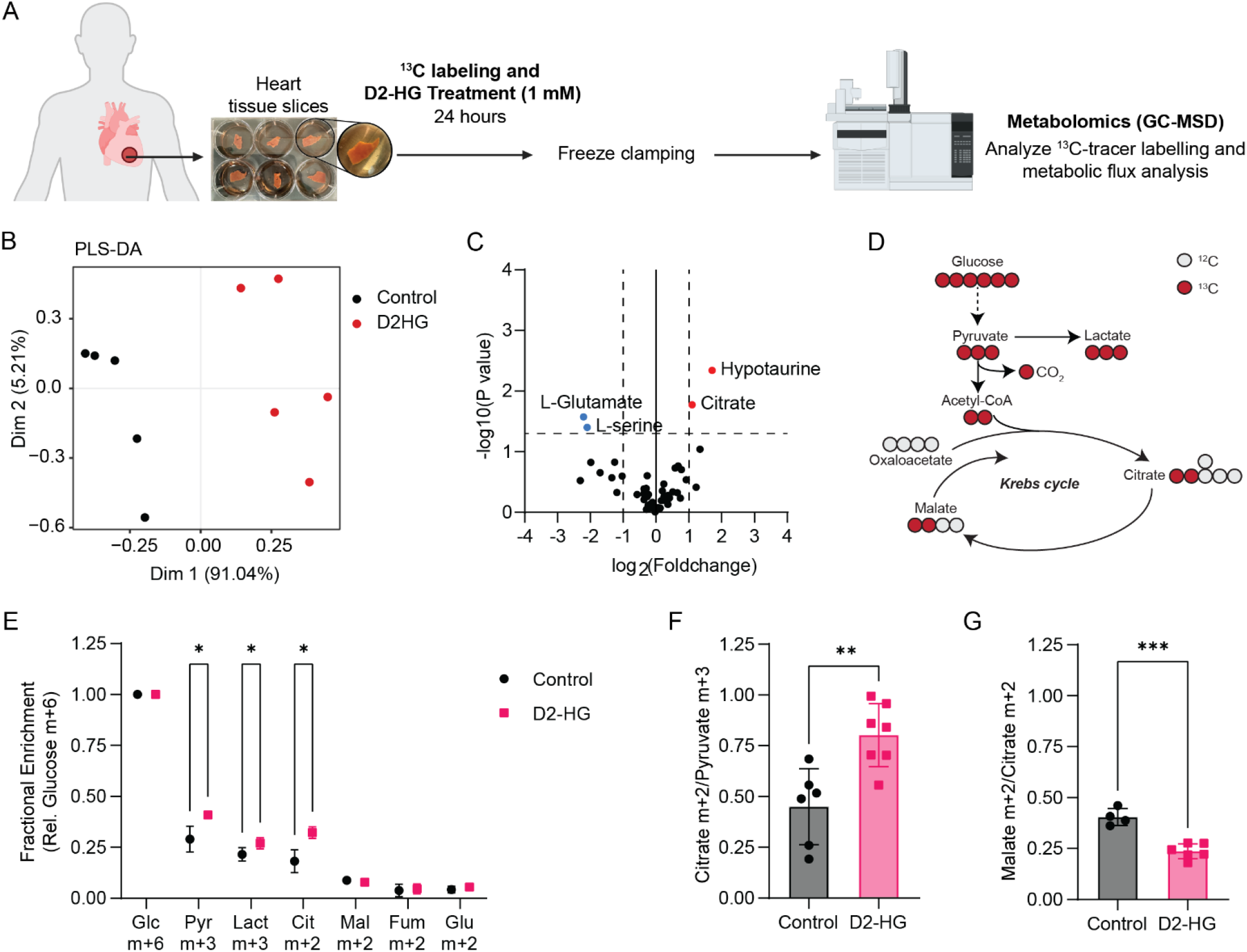
Glucose metabolism in human hearts during oncometabolic stress by D2-HG. (**A**) Schematic of donor heart collection. 500-μm sections of left ventricular heart tissue samples from 6-7 donors were treated with or without D2-HG, followed by *in vitro* ^13^C-labeling and gas chromatography and mass spectrometry (GC-MS)-based metabolomics. Tissue slices from the same patient served as both the control and treatment groups. (**B-C**) Partial Least Squares Discriminant Analysis (PLS-DA) (**B**) and volcano plot analysis (**C**) of metabolic profiles from human heart tissue slices treated with or without D2-HG (1.0 mM). (**D**) Isotopologue labeling of Krebs cycle intermediates via [U-^13^C]glucose. Colored circles indicate labeled carbons from [U-^13^C]-glucose. (**E**) ^13^C enrichment comparisons derived from glucose m+6 between human heart tissue slices treated with and without D2-HG (1.0 mM). The isotopologue of each metabolite is normalized to glucose m+6. Statistical significance was assessed using a Mann-Whitney test with a false discovery rate (FDR) and a two-stage step-up method of Benjamini, Krieger, and Yekutieli (FDR cutoff < 0.01). *p-value<0.05, **p-value<0.01,****p-value<0.001. (**F-G**) Citrate m+2 to pyruvate m+3 ratio (**F**) and malate m+2 to citrate m+2 ratio (**G**) from in vitro [U-^13^C]glucose labeling. Each point reflects one cultured human tissue slice. Statistical significance was assessed using a Mann-Whitney test with a false discovery rate (FDR) and a two-stage step-up method of Benjamini, Krieger, and Yekutieli (FDR cutoff < 0.01). **p-value<0.01.

### Glucose and glutamine feed the Krebs cycle during oncometabolic stress in the heart

To assess the acute metabolic consequences of D2-HG, we used the working rodent heart preparation as a model. We perfused wild-type (WT) rat hearts with or without D2-HG (1.0 mM) for 30 min under physiological nutrient concentrations and a normal workload. Consistent with our previous reports, cardiac hydraulic power and myocardial oxygen consumption were significantly reduced with D2-HG treatment compared to control conditions (**Supplementary Figure 2A** and **Figure 2A**). We perfused rat hearts with [U-^13^C]glucose to assess the fates of carbon from glucose-dependent pathways, including glycolysis and the Krebs cycle (**Figure 2B**). D2-HG levels were increased in perfused hearts (**Supplementary Figure 2B**). Analysis of total metabolite abundances using GC-MS revealed an increased accumulation of glutamine and glutamate (**Supplementary Figure 2C**). Consistent with human heart studies, perfused rat hearts with D2-HG displayed increased ^13^C labeling in pyruvate compared to control (**Figure 2C**). M+2 labeling in the Krebs cycle intermediates relative to pyruvate m+3 can be used as a surrogate for carbon entry through pyruvate dehydrogenase (PDH). On the other hand, m+3 labeling of Krebs cycle intermediates relative to pyruvate m+3 can be used as a surrogate for carbon entry via pyruvate carboxylase (PC) (**Figure 2B**). We observed increased transfer of glucose-derived pyruvate into the Krebs cycle via PDH in hearts perfused with D2-HG, resulting in an increased citrate m+2-to-pyruvate m+3 ratio (**Figure 2D**). In contrast, carbon transfer via PC was reduced, resulting in a decreased m+3 labeling in aspartate, malate, and citrate (**Figure 2E**). The citrate isotopologue distribution revealed decreased citrate m+3 and m+4 fractions in hearts perfused with D2-HG compared to controls (**Figure 2F**), suggesting reduced Krebs cycle flux downstream of citrate and reduced carbon incorporation at subsequent cycle turns. Next, we conducted metabolic flux analysis (MFA) using INCA^29^. Isotopologues were incorporated into a network that defines carbon transitions, and labeling patterns were fitted to the model. MFA revealed increased glycolytic flux and reduced Krebs cycle flux downstream of citrate (**Figure 2G**).

**Figure 2.**
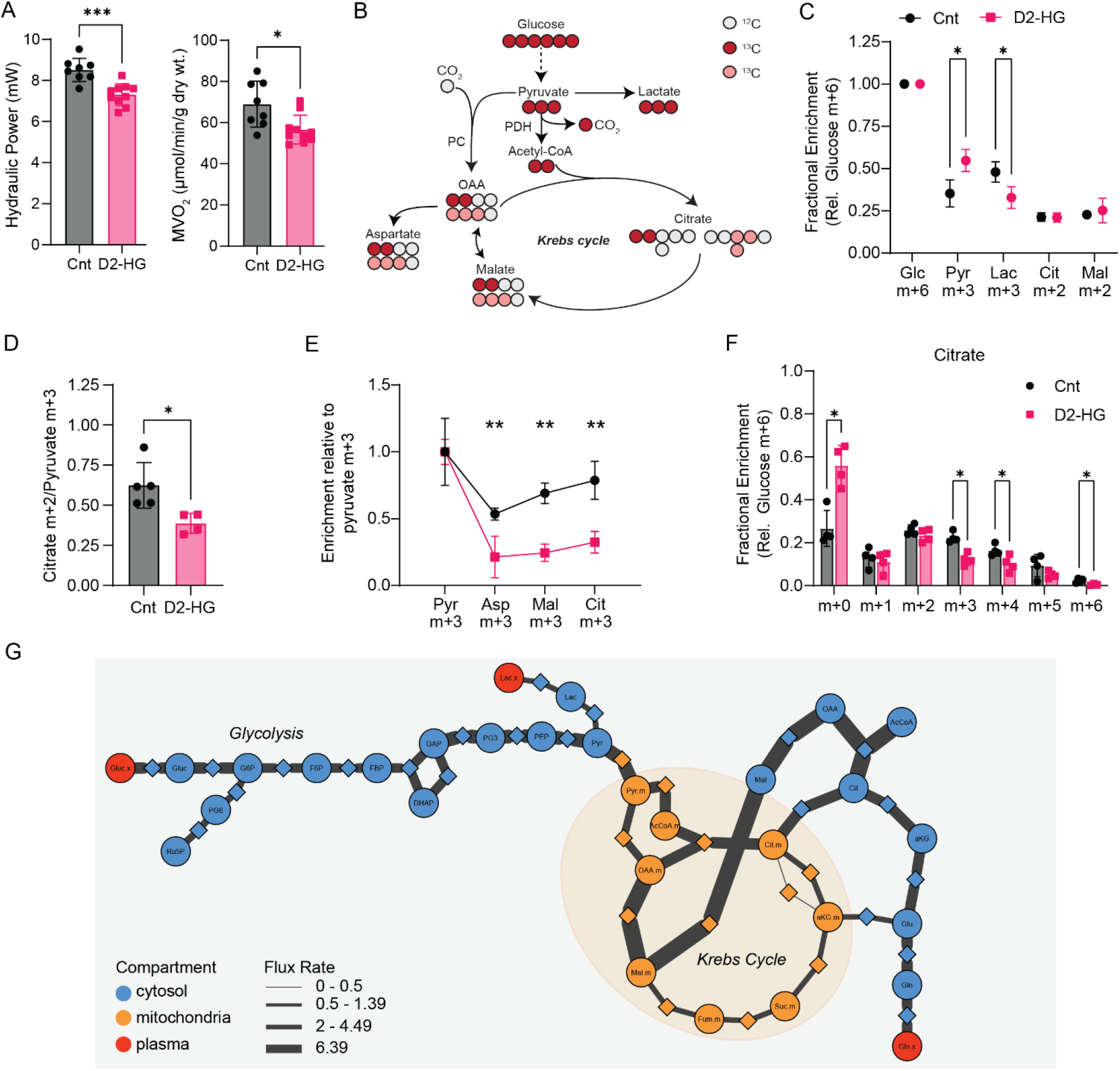
Reductive formation of citrate in the heart during oncometabolic stress by D2-HG. (**A**) Hydraulic power and myocardial oxygen consumption in ex vivo perfused working rat hearts with or without D2-HG (1.0 mM). N=8-10 male rats/group. Statistical analysis was conducted using the Mann-Whitney U test. ****p-value<0.001. (**B**) Schematic of carbon transitions from [U-^13^C]-glucose. Colored circles indicate labeled carbons from [U-^13^C]-glucose. Abbreviations: OAA, oxaloacetate; PDH, pyruvate dehydrogenase; PC, pyruvate carboxylase. (**C**) ^13^C enrichment comparisons derived from glucose m+6 between rat hearts perfused with and without D2-HG (1.0 mM). The isotopologue of each metabolite is normalized to glucose m+6. Statistical analysis was conducted using the Welch t-test with FDR<5%. *p-value<0.05, **p-value<0.01,****p-value<0.001. (**D**) Citrate m+2 to pyruvate m+2 ratio from ex vivo [U-^13^C]glucose labeling. Statistical analysis was conducted using an unpaired two-tailed t-test with Welch’s correction. N=8-10 male rats/group. **p-value<0.01. (**E**) Comparison of PC flux derived from pyruvate m+3. The isotopologs of each metabolite were normalized to pyruvate m+3. Statistical analysis was conducted using the Mann-Whitney U test with FDR<5%. *p-value<0.05, **p-value<0.01,****p-value<0.001. (**F**) Mass isotopomer analysis of citrate in perfused working rat hearts with [U-^13^C]-glucose. Statistical analysis was conducted using the Mann-Whitney U test with FDR<5%. *p-value<0.05, **p-value<0.01,****p-value<0.001. (**G**) Comparative metabolic flux analysis using ^13^C_6_-glucose isotopomer distributions from rat hearts perfused with or without D2-HG (1.0 mM). Carbon transitions were assigned to cytosolic, mitochondrial, or extracellular metabolic fluxes. The thickness of each line connecting metabolites and proteins indicates the calculated flux rates.

### Alternative Energy Sources during Oncometabolic Stress

The elevation of glutamate and the suppression of citrate-to-malate conversion in the Krebs cycle suggested that alternative carbon sources to glucose contribute to the production of Krebs cycle intermediates in the heart during D2-HG-producing IDH-mutant tumors. We perfused rat hearts for 30 min at physiological nutrient concentrations (see Methods), with or without D2-HG and ^13^C tracers. We first tested if D2-HG is a carbon donor in perfused rat hearts using [U-^13^C]-D2-HG. Enrichment of D2-HG m+5 was readily achieved within the duration of the perfusion protocol (**Supplementary Figure 3A**). D2-HG can be converted to α-KG via D-2-hydroxyglutarate dehydrogenase (D2HGDH), thereby supporting oxidative metabolism via anaplerosis of α-KG rather than glutamine. Surprisingly, we did not observe a substantial incorporation of D2-HG-derived carbons into Krebs cycle intermediates (**Figure 3A** and **Supplementary Figure 3B-H**). The contribution of D2-HG to glutamine m+5 and citrate m+5 was 2.39±0.29% and 0.219±0.04%, respectively. Our data indicate that, during the perfusion protocol, D2-HG contributed only minimally to oxidative metabolism. Next, we perfused hearts with [U-^13^C]glutamine to determine the incorporation of glutamine into Krebs cycle intermediates. Glutamine contributes to oxidative and reductive pathways (**Figure 3B**), which contribute ^13^C to the Krebs cycle via α-KG. Oxidative decarboxylation of α-KG via α-KGDH leads to the formation of glutamine-derived citrate m+4 in the first turn (**Figure 3B**). In contrast, the reductive pathway converts glutamine-derived α-KG to citrate m+5 via cytosolic IDH1 or mitochondrial IDH2 (**Figure 3B**). We observed increased citrate m+5 from [U-^13^C]glutamine labeling (**Figure 3C**), consistent with increased reductive formation of citrate from exogenous glutamine. Malate m+3 downstream of citrate also increased (**Figure 3C**). To further differentiate between reductive and oxidative citrate generated from glutamine, we next perfused rat hearts with [1-^13^C]glutamine. Oxidative conversion of glutamine to citrate will remove only the labeled carbon and yield unlabeled citrate (m+0), while reductive conversion will yield citrate m+1 (**Figure 3B**). We observed an increased glutamine-derived citrate m+1 with D2-HG perfusions compared to control perfused rat hearts (25.33±8.5% vs. 4.4%±1.5%, q-value=0.008; **Figure 3D**), indicating that during oncometabolic stress, the reductive formation of citrate is a significant contributor to metabolic adaptation and compensation in the heart. Reductive and oxidative metabolism require the regeneration of redox equivalents in the form of NADH and NADPH. We next quantified the contribution of glucose to NADH regeneration via glycolysis using [3-^2^H]glucose in perfused working rat hearts. In the glycolysis, glyceraldehyde 3-phosphate dehydrogenase (G3PDH) converts glyceraldehyde 3-phosphate (G3P) to 1,3-bisphosphoglycerate (1,3BPG) and simultaneously reduces NAD^+^ to NADH, which gains a ^2^H-labelled molecule (**Figure 3E**). Further reduction of 1,3BPG leads to the loss of ^2^H within the glycolysis. Therefore, [3-^2^H]glucose allows to determine the glucose-derived contribution of NADH. Perfusions were conducted at normal workload for 30 min with physiological concentrations of nutrients (See Methods) and with or without D2-HG (1.0 mM). MS-based metabolomics revealed a reduced abundance of NADH (**Figure 3F**) in rat hearts perfused with D2-HG compared to controls. The fractional enrichment of ^2^H-NADH was significantly reduced in D2-HG-perfused rat hearts compared to controls, indicating that glucose-derived ^2^H is directed at the level of glucose 6-phosphate into the pentose phosphate pathway to generate ^2^H-NADPH. Based on these results, we reasoned that either NADPH-dependent cytosolic IDH1 or mitochondrial IDH2 catalyzes the reductive formation of citrate.

**Figure 3.**
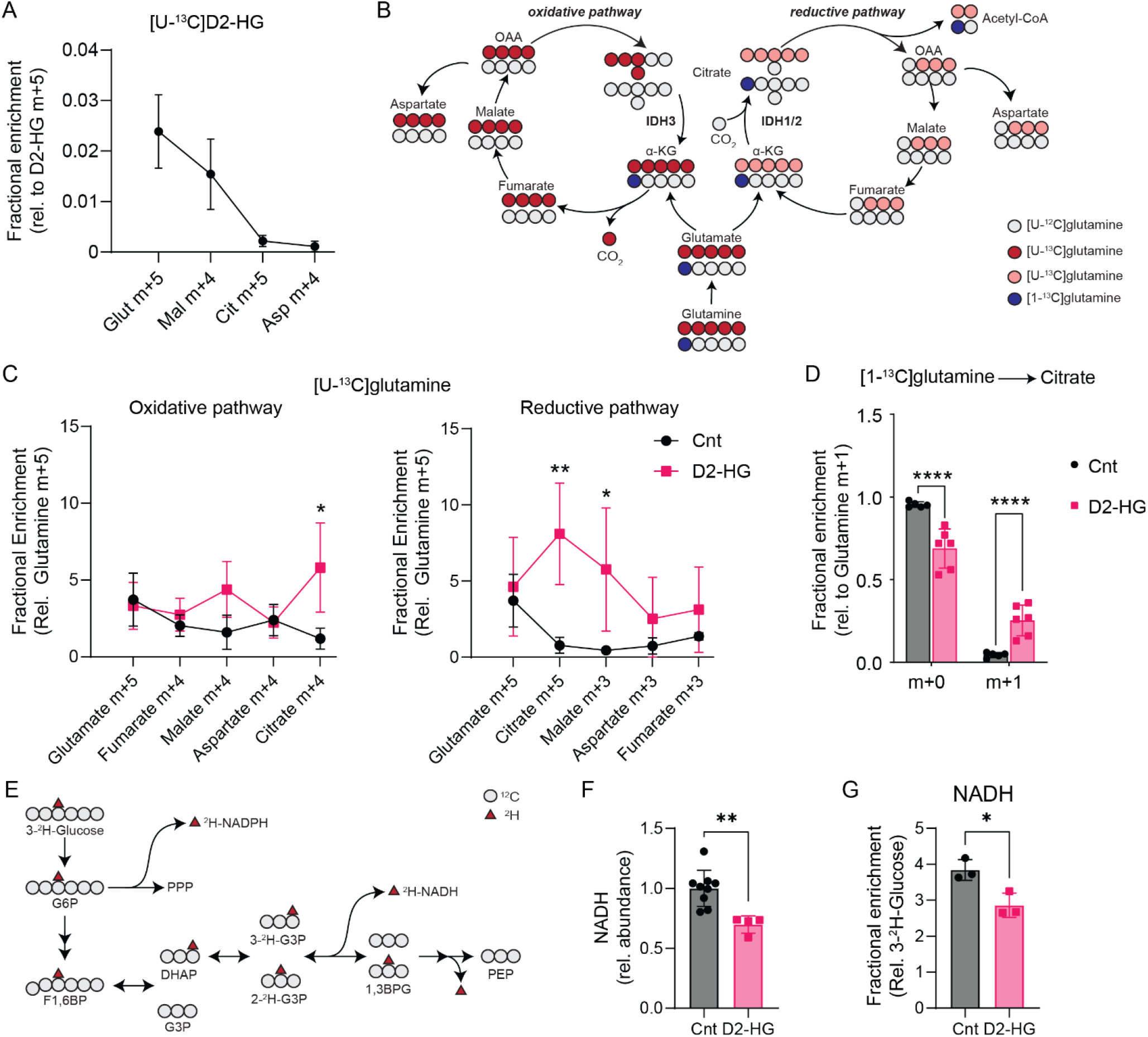
Glutamine is a major citrate precursor in hearts perfused with D2-HG. (**A**) Contribution of [U-^13^C]-D2-HG to Krebs cycle intermediates in rat hearts perfused with D2-HG (1 mM). The isotopologue of each metabolite is normalized to D2-HG m+5. (**B**) Schematic of reductive and oxidative carbon transitions from [U-^13^C]-glutamine. Colored circles indicate labeled carbons from [U-^13^C]-glutamine (red) or [1-^13^C]- glutamine (blue). Abbreviations: IDH, isocitrate dehydrogenase; OAA, oxaloacetate. (**C**) Mass isotopologue analysis of Krebs cycle intermediates representing oxidative (left panel) and reductive (right panel) pathways in rat hearts perfused with [U-^13^C]-glutamine and unlabeled glucose. Statistical analysis was conducted using the Mann-Whitney U test with FDR<5%. *p-value<0.05, **p-value<0.01. (**D**) Fractional enrichment of citrate from [1-^13^C]-glutamine in rat hearts perfused with or without D2-HG (1 mM). Statistical analysis was conducted using the Mann-Whitney U test with FDR<5%. ****p-value<0.001. (**E**) Schematic of [3-^2^H]- glucose transition to ^2^H-NADH from glycolytic flux. Colored triangles indicate labeled hydrogen from [3-^2^H]-glucose. Abbreviations: G6P, glucose 6-phosphate; F1,6BP, fructose 1,6-bisphoshate; DHAP, dihydroxyacetone phosphate; G3P, glycerol 3-phosphate; 1,3BPG, 1,3-bisphosphoglyceric acid; PEP, phosphoenolpyruvate. (**F**, **G**) MS-based abundance of NADH (**F**) and fractional enrichment of [3-^2^H]-glucose-derived NADH (**G**) in rat hearts perfused with or without D2-HG (1 mM). Statistical analysis was conducted using the Mann-Whitney U test with FDR<5%. ****p-value<0.001.

### IDH1 drives reductive formation of citrate during oncometabolic stress

To test which IDH isoform drives reductive citrate formation in the heart, we used small interfering RNA (siRNA) to knock down IDH1, IDH2 or IDH3 in a human cardiomyocyte-derived cell line (AC16) (**Figure 4A**). Differentiated AC16 were transfected with non-targeting siRNA (wildtype control) or siRNA targeting IDH isoforms. We confirmed successful knockdown of IDH-isoform gene expression by RT-qPCR (**Supplementary Figure 4A)** and protein expression by Western blot (**Supplementary Figure 4B-D**). Transfected differentiated AC16 were cultured for 24h with or without D2-HG (1.0 mM) and labeled with ^13^C-tracers to elucidate the metabolic fates of glucose and glutamine. We found a marked increase in citrate m+1 in wildtype (WT) AC16 treated with D2-HG compared to controls (58%±17.2%, **Figure 4B**). Only the knockdown of IDH1 abrogated the reductive formation of citrate from [1-^13^C]glutamine (**Figure 4B**). Likewise, [U-^13^C]-glucose labeling demonstrated reduced citrate m+5 enrichment (**Figure 4C**) in AC16 treated with siRNA-IDH1, consistent with a decreased reductive formation of citrate. To corroborate these findings, we generated viable agarose-embedded heart tissue slices from three healthy human heart donors (**Figure 4D**). Heart tissue slices were randomly assigned to control or experimental groups, with tissue samples from the same patient serving as controls (see **Methods**). Tissue slices were treated with or without D2-HG (1.0 mM), the IDH1^R132H^-mutant enzyme inhibitor vorasidenib (5 μM; AG-120, IDH1i), or the IDH2^R140Q^-mutant enzyme inhibitor enasidenib (5 μM; AG-221, IDH2i). AG-120 and AG-221 display micromolar potency in inhibiting the IDH1^WT^ and IDH2^WT^ enzymes in cell lines^30^. To assess the potency of IDH inhibitors in human hearts, we used drug-affinity responsive target stability (DARTS), an unbiased molecular approach that quantifies the affinity of small-molecule targets to proteins. We found decreased proteinase K susceptibility of IDH1 in the presence of AG-120 (IDH1i) and AG-221 (IDH2i) (**Figure 4E**). This decreased susceptibility suggests that IDH inhibitors bind to and stabilize IDH1, affecting its activity. Tracing with [1-^13^C]-glutamine revealed a markedly increased labeling of citrate m+1 in the presence of D2-HG compared to control (82.95±14.8% vs. 14.87±7.1%, **Figure 4F**). Treatment with AG-120 (IDH1i) abrogated citrate m+1 enrichment in D2-HG (28.8±10.67%, **Figure 4F**), while AG-221 (IDH2i) showed a downward trend yet no significant difference compared to D2-HG-treated tissue slices (61.1±5.37%, **Figure 4F**). Together, our data demonstrate that IDH1 is the route of citrate synthesis from glutamine during oncometabolic stress in human hearts.

**Figure 4.**
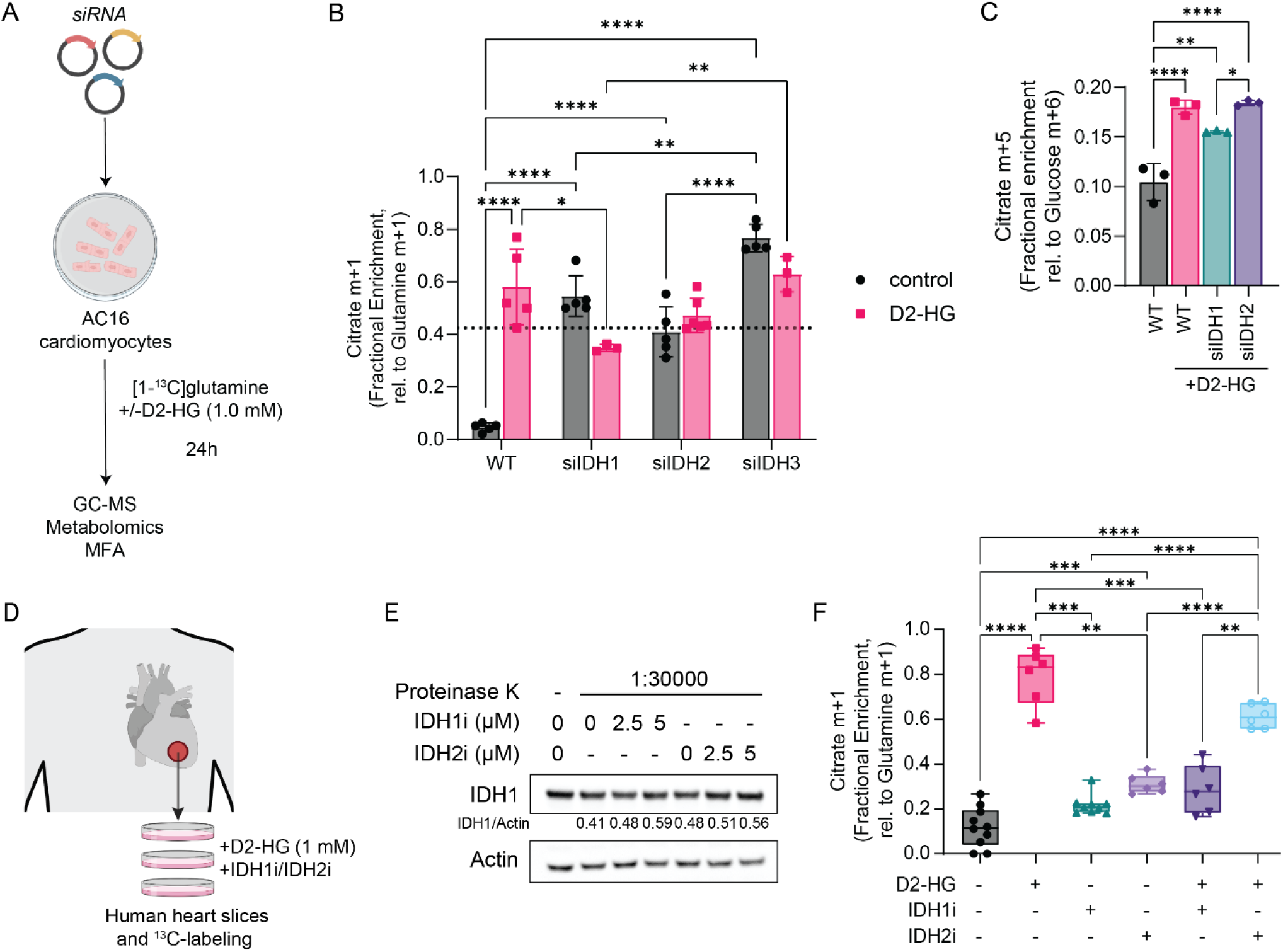
Inhibition of NADPH-dependent IDH isoforms attenuates reductive carboxylation in human heart tissue slices. (**A**) Transient silencing of isocitrate dehydrogenase (IDH) in cultured differentiated human AC16 cardiomyocytes with short interfering RNAs (siRNA) directed against IDH1, IDH2, or IDH3. A non-targeting siRNA was used as a negative control (WT). (**B, C**) Mass isotopomer analysis of citrate in differentiated human AC16 cardiomyocytes with [1-^13^C]-glutamine (**B**) and [U-^13^C]-glucose (**C**) after silencing IDH isoforms. Statistical analysis was conducted using 2-way ANOVA followed by multiple comparisons analysis with FDR<5%. *p-value<0.05, **p-value<0.01, ***p-value<0.005, ****p-value<0.001. (**D**) Workflow of human donor heart tissue slices and ^13^C-tracer labeling. (**E**) DARTs blotting showing AG-120 (IDH1 inhibitor, IDH1i) and AG-221 (IDH2 inhibitor, IDH2i) as a substrate of IDH1. The susceptibility of IDH1 to proteinase K digestion is increased in the presence of AG-120 and AG221. (**F**) Mass isotopomer analysis of citrate in cultured human heart tissue slices with [1-^13^C]-glutamine after treatment with AG-120 (IDH1 inhibitor, IDH1i) and AG-221 (IDH2 inhibitor, IDH2i). Statistical analysis was conducted using Brown-Forsythe and Welch ANOVA tests and multiple comparisons analysis with Dunnetts T3. *p-value<0.05, **p-value<0.01, ***p-value<0.005, ****p-value<0.001.

## DISCUSSION

We have previously demonstrated that D2-HG impairs cardiac contractile function in rodent hearts through the inhibition of α-KGDH^9^. We have demonstrated that D2-HG causes reductive carboxylation of α-KG to citrate via IDH1 in human and rodent hearts. NADPH-dependent IDH1 and IDH2 are reversible, which enables the reductive formation or oxidative decarboxylation of citrate. We identified IDH1 as the critical driver for reductive carboxylation in the heart using three models: *in vitro* AC16 cardiomyocytes, *ex vivo* working rat heart perfusions, and human heart tissue slices. Across all models, we observed that D2-HG promotes redirection of Krebs cycle intermediates and increases reductive carboxylation via IDH1, which is abrogated by inhibiting IDH1.

D2-HG increases glutamine metabolism in the perfused rodent heart and human heart tissue samples. The increased contribution of glutamine to oxidative and reductive metabolism is a hallmark of metabolic stress^31^. Interestingly, D2-HG does not transfer carbon to Krebs cycle intermediates; instead, it shifts nutrient contributions through allosteric inhibition of α-KGDH. Our tracer labeling data from D2-HG-treated human and rodent hearts are similar to persistent glutamine oxidation observed in cancer cells^18,22,32^ and during ischemia-reperfusion injury in the heart^33^. Glutamine is not a primary energy-providing substrate in the heart, as supported by the limited contribution of citrate m+4 during rodent heart perfusion with [U-^13^C]glutamine. A lack of glutamine oxidation is consistent with an impaired Krebs cycle function, as supported by the limited glucose-derived ^13^C-citrate labeling and enrichment in m+3 and m+4, suggesting incomplete Krebs cycle turnover. Together, these factors are predicted to increase the reductive formation of citrate from glutamine, which we observed as citrate m+5.

Knockdown of IDH isoforms in vitro using AC16 cardiomyocytes revealed that only IDH1 attenuated reductive citrate formation, confirming that this IDH isoform carries reductive flux in response to oncometabolic stress. Intriguingly, even in the absence of D2-HG, the knockdown of IDH3 increased reductive citrate formation. Loss of IDH3 expression has been demonstrated in advanced heart failure and preclinical models^34–36^. Our data suggest that loss of IDH3 expression before oncometabolic stress will increase reductive metabolism and further exacerbate mitochondrial dysfunction in the heart. These findings are surprising because IDH2 activity could compensate for IDH3 loss. Our data suggest that limited mitochondrial NADPH supply may prevent compensatory IDH2 flux and raise the possibility of increased reductive citrate formation during HF, independent of D2-HG-producing tumors.

NADH and NADPH synthesis play critical roles in regulating the progression of cardiac remodeling and HF^37,38^. Increased glucose uptake and conversion to glucose 6-phosphate promote flux into the oxidative branch of the pentose phosphate pathway, which synthesizes NADPH. Therefore, IDH1 and IDH2 functions are directly linked to glucose metabolism via the regeneration of NADH and NADPH. Tracer labeling with [3-^2^H]glucose showed decreased NADH synthesis through glycolysis, suggesting that glucose 6-phosphate is redirected into the pentose phosphate pathway, contributing to NADPH synthesis. This function supports NADPH-consuming IDH1 flux and reductive formation of citrate from glutamine in the cytosol. We observed increased citrate levels in human heart tissue and cultured cardiomyocytes. Citrate is an allosteric inhibitor of phosphofructokinase downstream of glucose 6-phosphate, which is diverted into the pentose phosphate pathway, providing precursors for NADPH synthesis^39^. This redirection of glycolytic intermediates promotes substrate channeling and positive feedback for IDH1 flux. Likewise, reduced regeneration of NAD^+^ to NADH in glycolysis supports channeling of NAD^+^ to fatty acid oxidation and autophagic adaptation^10^. Therefore, a possible explanation for the reductive formation of citrate in the cytosol is that while IDH2 enhances the conversion of Krebs cycle intermediates in the mitochondria, the physical relationship with glucose metabolism supports NADPH and precursor synthesis.

The importance of metabolic adaptation as a therapeutic target in the heart is becoming increasingly apparent. Efforts to improve cardiac function by suppressing cardiac metabolism require understanding how energy-providing substrates are redirected into other pathways during periods of stress and how compensatory fluxes are regulated. Our study provides evidence for a synergistic regulation of oxidative metabolism via cytosolic IDH1, but a challenge remains in determining how these changes affect structural remodeling in the heart. Pressure-overload-induced heart failure mouse models have demonstrated reduced gene expression of Krebs cycle enzymes, including IDH3^34,36,40^. It seems likely that if mitochondrial impairment induces compensatory IDH1 flux, targeting IDH1 has therapeutic benefits.

A limitation of the tracer studies described so far, in human heart tissue slices and cultured cardiomyocytes, is that these experiments may overestimate the contribution of glutamine. Recent work in cell culture models of cancer demonstrates that culture conditions increase the metabolic flux from glutamine^40,41^. Nevertheless, the current work provides evidence that oncometabolic stress increases the reductive formation of citrate via IDH1 and that this activity supports glutamine uptake. The reduced citrate m+1 labeling from glutamine following IDH1 knockdown or pharmacologic inhibition demonstrates that IDH1 is a targetable metabolic vulnerability in the human heart. Our studies support the development of therapeutic strategies that target cancer cell metabolism and oncometabolic remodeling of the heart.

## ACKNOWLEDGEMENTS

Figures were created with BioRender.com and Adobe Illustrator.

## AUTHOR CONTRIBUTIONS

Conceptualization, A.K.; Methodology, K.K. and A.K.; Metabolomics, A.K., K.K., N.S., B.F., R.D.; Computational Modeling, A.K., K.K.; Data analysis, K.K., Y.G., A.K.; Investigation, Y.G., K.K., I.W., E.V.; Writing – Original Draft, K.K. and A.K.; Writing – Review & Editing, Y.G., K.K., I.W., N.S., E.V., R.D., B.F., H.T.; Resources, H.T., R.D.

