## Supplemental_Materials for "Reductive carboxylation via isocitrate dehydrogenase 1 supports cardiac metabolic adaptation during oncometabolic stress"

**TITLE**

**SHORT-TITLE**

IDH1 Causes Reductive Carboxylation in the Heart

**CORRESPONDING AUTHOR**

Anja Karlstaedt, MD, PhD
Assistant Professor
Department of Cardiology, Smidt Heart Institute
Cedars Sinai Medical Center
127 San Vicente Blvd, Advanced Health Science Pavilion 9229
Los Angeles, California 90048, USA


### METHODS

#### Animal Care and Studies

All animal experiments were conducted on male rats, approved by the Institutional Animal Care and Use Committee at Cedars Sinai Medical School, and carried out in accordance with the National Institute of Health guidelines. Wildtype rats were purchased from Charles River. Only male rats were used in the study. All *ex vivo* experiments were performed by IACUC approval at Cedars Sinai Medical Center (Los Angeles, CA, USA) and The University of Texas Health Science Center (Houston, TX, USA), MD Anderson Cancer Center (Houston, TX, USA). Rats had free access to water and food, which consisted of standard chow. Animals were housed in colony cages (maximum of 2 animals per cage) with 12-hour shifts of the light-dark cycle. Data management and productivity planning of the rats was facilitated by Transnetyx Colony Management and Scinote Electronic Lab Notebook Software.

#### Working Rodent Heart Perfusion

Rat hearts were perfused by the method described earlier.[1] Briefly, rats were anesthetized with pentobarbital (100 mg/kg, intraperitoneal) and heparinized (200 U) through direct injection into the inferior *V. cava* after laparotomy. Next, the chest was opened, and the heart rapidly excised and arrested in ice-cold Krebs-Henseleit (KH) buffer (120 mmol/L NaCl, 5 mmol/L KCl, 1.2 mmol/L $MgSO_4$, 1.2 mmol/L $KH_2PO_4$, 25 mmol/L $NaHCO_3$, 2.5 mmol/L $Ca^{2+}$) at pH 7.4. Hearts were perfused in the working mode for 30 min with Krebs-Henseleit (KH) buffer containing physiologic levels of substrates, including 5 mM [U-$^{13}$C]-glucose, 0.5 mM L-lactate, 1 mM glutamine, and 0.1 mM palmitate bound to albumin, as described previously. [2,3] For treatments, D-2-hydroxyglutarate (D2-HG, 1.0 mmol/L; Tocris Bioscience, Minneapolis, MN, USA; Cat. No. 6124) was added to the perfusion buffer. For labeling experiments, glucose and glutamine were replaced with stable isotope tracers (Cambridge Isotope Laboratories, Inc., Boston, MA): [U-$^{13}$C]glucose (CLM1396-5), [U-$^{13}$C]glutamine (CLM-1822-H-PK) or [1-$^{13}$C]glutamine (CLM-3612-PK), respectively. At the end of the perfusion protocol, hearts were freeze-clamped using aluminum tongs cooled in liquid nitrogen and stored at -80°C for further tissue analysis.

#### AC16 Cell Culture.

The AC16 human ventricular cardiomyocyte line (Cat.No.#CRL-3568, ATCC, Manassas, VA) was maintained in a complete DMEM medium (Cat. No. 10-017-CV, Corning, Thermo Fisher Scientific, Carlsbad, CA) at 37°C and 5% $CO_2$. Cells were differentiated for 72h using DMEM:F-12 media (Cat. No. 31331018, Gibco, Thermo Fisher Scientific, Waltham, MA) complemented with Insulin-Transferrin-Selenium (Cat. No. 41400045, Gibco, Thermo Fisher Scientific, Waltham, MA) and 2% Serum Replacement II (Cat. No. 14009C-500ML, Sigma Aldrich, St. Louis, MO, USA). Differentiated AC16 cells were cultured in DMEM without phenol red supplemented with glucose (5.5mM), glutamine (0.1 mM), and pyruvate (0.5 mM) at 37°C and 5% $CO_2$. Cells were treated for 24h with phosphate-buffered saline (PBS, Sigma Aldrich, St. Louis, MO, USA; Cat. No. 506552) or D-2-hydroxyglutarate (D2-HG, 1.0 mmol/L; Tocris Bioscience, Minneapolis, MN, USA; CAT#6124). For labeling experiments, glucose and glutamine were replaced with stable isotope tracers (Cambridge Isotope Laboratories, Inc., Boston, MA): [U-$^{13}$C]glucose (Cat. No. CLM1396-5), [U-$^{13}$C]glutamine (Cat. No. CLM-1822-H-PK) or [1-$^{13}$C]glutamine (Cat. No. CLM-3612-PK), respectively.

#### IDH silencing in AC16

Knockdown of IDH isoforms was achieved *in vitro* using silencing RNA (siRNA) (1µM, Horizon Dharmacon, Lafayette, CO, USA): IDH1 (NM_005896, Cat. No. E-008294-00-0010), IDH2 (NM_002168, Cat. No. E-004013-00-0010), IDH3 (NM_174856, Cat. No. E-008753-00-0010) and non-targeting siRNA (Cat. No. D-001910-10-05). Differentiated AC16 were transfected for 72h in serum-free cell culture media using predesigned Accell siRNA without transfection reagent per the manufacturer's instructions. Following transfection, differentiated AC16 were cultured with cultured in DMEM without phenol red supplemented with glucose (5.5 mM), glutamine (0.1 mM), and pyruvate (0.5 mM) at 37°C and 5% $CO_2$. Cells were treated for 24h with phosphate-buffered saline (PBS, Sigma Aldrich, St. Louis, MO, USA; Cat. No. 506552) or D-2-hydroxyglutarate (D2-HG, 1.0 mmol/L; Tocris Bioscience, Minneapolis, MN, USA; Cat. No. 6124). For labeling experiments, glucose and glutamine were replaced with stable isotope tracers (Cambridge Isotope Laboratories, Inc., Boston, MA): [U-$^{13}$C]glucose (Cat. No. CLM1396-5) or [1-$^{13}$C]glutamine (Cat. No. CLM-3612-PK), respectively.

**Human Heart Tissue Slices**

Adult human hearts were procured as donor human hearts from the UNOS Transplant Services. Experimental protocols were approved by the Smidt Heart Institute Tissue Repository (Protocols: CS-IRB Pro00010979 and Pro00011910). Informed consent was obtained from all patients. Human heart tissue samples were obtained from donor patients, which were rejected for transplantation due to organ size, lack of organ recipient, or ischemia time. Explanted hearts were cardioplegically arrested via high potassium solution (in mmol/L: NaCl 110, $CaCl_2$ 1.2, KCl 16, $MgCl_2$ 16, $NaHCO_3$ 10; Sigma Aldrich, St. Louis, MO, USA) and were cooled to 4 °C in the operating room following aortic cross-clamp. Left ventricular tissue was dissected from regions near the left anterior descending and circumflex artery. Tissue samples were embedded in 1% agarose (Cat. No. 16520050, Thermo Fisher Scientific, Carlsbad, CA) and cut tangential to the endocardium at 0.5 µM. Tissue slices were immediately transferred into a 6-well cell culture plate using sterile forceps and gradually warmed to 37°C for tracer labeling and inhibitor studies. Slices were cultured in DMEM (Cat. No. A14430-01, Thermo Fisher Scientific, Carlsbad, CA) supplemented with glucose (5.5 mM), glutamine (0.1 mM), and pyruvate (0.1 mM). For oncometabolic studies, PBS (Cat. No. 506552, Sigma Aldrich, St. Louis, MO, USA) or D2-HG (1.0 mmol/L; Cat. No. 6124, Tocris Bioscience, Minneapolis, MN, USA) was added to the media. For labeling experiments, glucose and glutamine were replaced by [U-$^{13}$C]glucose (5.5 mM, Cat. No. CLM1396-5; Cambridge Isotope Laboratories, Inc., Boston, MA, USA) and [1-$^{13}$C]glutamine (0.1 mM, Cat. No. CLM-3612-PK; Cambridge Isotope Laboratories, Inc., Boston, MA, USA), respectively. For IDH1 and IDH2 inhibitor studies, ivosidenib (5µM, AG-120, Cat. No. HY-18767, MedChemExpress, Monmouth Junction, NJ, USA) and enasidenib (5µM, AG-221, Cat.No. HY-18690, MedChemExpress, Monmouth Junction, NJ, USA).

**Western Blotting**

The expression of proteins was analyzed using Western blotting. Proteins were extracted from flash-frozen tissues as described previously.[2,3] Briefly, tissue samples were homogenized and lysed in RIPA buffer (10 mmol/L Tris at pH7.5, 1 mmol/L EDTA, 0.1% SDS, 1% Triton X-100, 0.1% Sodium deoxycholate, 5 mol/L NaCl) containing EDTA-free Protease and Phosphatase inhibitors (Cat. No. #11873580001, Rose Holding AG, Basel, Switzerland). Proteins were separated on 4-12% SDS-PAGE gels (Precast Protein gels, Cat. No. NP0322BOX, Invitrogen, Waltham, MA, USA), transferred to PVDF membranes and probed with antibodies from Cell Signaling Technology (CS, Danvers, MA, USA) against IDH1 (Cat. No. 8137S, Cell Signaling Technology, Danvers, MA, USA), IDH2 (Cat. No. 56439S, Cell Signaling Technology, Danvers, MA, USA) and IDH3 (Cat. No. NBP2-1411, Novus International Inc., Chesterfield, MO, USA). Protein levels were detected by immunoblotting using horseradish peroxidase-conjugated secondary antibodies and chemiluminescence. Secondary antibodies: goat anti-rabbit IgG (Cat. No. 7074, Cell Signaling Technology, Danvers, MA, USA) and goat anti-rabbit IgG (Cat. No. 7076, Cell Signaling Technology, Danvers, MA, USA).

**Quantitative Real-Time-PCR Analysis**

Total RNA was extracted from AC16 cells using the RNeasy Plus Mini Kit for fibrous tissue (Cat#74034; Qiagen, Hilden, Germany) according to the manufacturer's instructions. RNA concentration was determined using a Qubit 4 Fluorometer (Cat. No. Q33226, Invitrogen, USA; RRID:SCR_018095). cDNA was synthesized using the iScript™ cDNA

Synthesis Kit according to the manufacturer's instructions (Cat. No. 1708890, Bio-Rad Laboratories, Hercules, CA). Quantitative real-time PCR was conducted using SYBR Green probes and measured using the QuantStudio using the following PCR settings: stage 1, 50°C 2 min, 95°C 10 min; stage 2, 95°C 15 sec, 60°C 1 min, cycles 40. Results were analyzed as ΔΔCt and expressed as the fold change in transcript levels. Reference gene expression was determined using the BioRAD reference gene panel. Expressions were quantified using PrimePCR SYBR Green Assay primers (Cat. No. 10025636, Bio-Rad Laboratories, Hercules, CA

**GC-MS Metabolomics.**

Tissue samples from rat heart perfusions were collected at the end of the perfusion protocol using freeze clamping. Human heart tissue samples were freeze-clamped after in vitro tracer labeling. Frozen tissue samples (5-10mg) were homogenized in MS-grade methanol, acetonitrile, and water at a volume ratio of 40:40:20. Homogenization of the tissue samples was achieved by freeze-thawing samples three times. Extracts were centrifuged at 13,000 g for 5 min at 4°C. Then, the supernatant was transferred to clean glass tubes. Myristic acid D27 (Cat.No. 366889; Sigma-Aldrich, St. Louis, MO) was used as an internal standard. The supernatant was transferred to clean glass tubes, and D27-myristic acid (0.15 µg/µL, Cat. No. 366889, Sigma- Aldrich; St Louis, MO) was added as internal run standard. Vials were covered with a breathable membrane, and metabolites were evaporated to dryness under vacuum and stored at -80°C for further GC-MS analysis. For derivatization of polar metabolites, frozen and dried metabolites were dissolved in 20 µL of Methoxamine (MOX) Reagent (2% solution of methoxyamine-hydrogen chloride in pyridine; Cat. No. 89803 and 270970; Sigma- Aldrich; St Louis, MO) and incubated at 30°C for 90 minutes. Subsequently, 90 µL of N-tert-Butyldimethylsilyl-N-methyltrifluoroacetamide with 1% tert-Butyldimethyl-chlorosilane (TBDMS; Cat. No. 375934, Sigma- Aldrich; St Louis, MO) was added, and samples were incubated under shaking at 37°C for 30 minutes. GC-MS analysis was conducted using an Agilent 8890 GC coupled with an Agilent 5977-mass selective detector. Metabolites were separated on an Agilent HP-5MS Ultra Inert capillary column (Cat. No. 19091S-433UI). For each sample, 1 µL was injected at 250°C using helium gas as a carrier with a flow rate of 1.1064 mL/min. For the measurement of polar metabolites, the GC oven temperature was kept at 60°C and increased to 325°C at a rate of 10°C/min (10 min hold), followed by a post-run temperature at 325°C for 1 min. The total run time was 37 min. The MS source and quadrupole were kept at 230°C and 150°C, respectively, and the detector was run in scanning mode, recording ion abundance within 50 to 650 m/z. **Supplemental Table 1** lists transitions and carbons for metabolites used in $^{13}$C-tracer analysis. Date were corrected for natural isotope effects using isotope correction software (IsoCor)[4].

| Metabolite | Carbons | Formula | m/z |
|---|---|---|---|
| Aspartate | 1,2,3,4 | C4H2NO4 | 418 |
| Citrate | 1,2,3,4,5,6 | C6H4O7 | 459 |
| Fumarate | 1,2,3,4 | C4H2O4 | 287 |
| Glutamate | 1,2,3,4,5 | C5H5NO4 | 432.3 |
| Alanine | 1,2,3 | C3H5NO2 | 160 |
| Glutamine | 1,2,3,4,5 | C5H6N2O3 | 431 |
| Pyruvate | 1,2,3 | C3H4O3 | 174 |
| Lactate | 1,2,3 | C3H4O3 | 261 |
| Malate | 1,2,3,4 | C4H2O5 | 419 |
| Serine | 1,2,3 | C3H4NO3 | 390 |

**Supplemental Table 1. Metabolite transitions used in metabolic flux analysis.**

**Metabolic Flux Analysis (MFA).**

Metabolic flux analysis was conducted using Isotopomer network compartmental analysis (INCA) platform[5] and MATLAB (v. 2023b, Optimization Toolbox and Statistics and Machine Learning Toolbox) by minimizing the sum of squared residuals (SSR) between model-simulated and experimental metabolite labeling measurements. Glutamate and glucose labeling from perfused working rat hearts or culture human heart tissue slices were provides as inputs to INCA simulations. Best-fit metabolic flux solutions were determined for each experiment by least-squares regression of the experimental measurements using the isotopomer network model (**Supplementary Table 2**). To ensure optimal solution was obtained, flux estimations were repeated a minimum of 100 times from randomized initial guesses with a relative convergence tolerance of 0.05. A chi-square test was used to assess goodness-of-fit, and a sensitivity analysis was conducted to determine 95% confidence intervals for each calculated flux.

| Reaction ID | Reaction | Carbon Transition |
|---|---|---|
| R1 | Gluc.x -> Gluc | abcdef -> abcdef |
| R2 | Lac -> Pyr | abc -> abc |
| R3 | Pyr -> Pyr.m | abc -> abc |
| R4 | Lac -> Lac.x | abc -> abc |
| R5 | Gluc -> G6P | abcdef -> abcdef |
| R6 | F6P -> FBP | abcdef -> abcdef |
| R7 | PEP -> Pyr | abc -> abc |
| R8 | G6P -> PG6 | abcdef -> abcdef |
| R9 | PG6 -> Ru5P + CO2 | abcdef -> abcde + f |
| R10 | Ru5P -> sink | abc -> |
| R11 | PG3 -> PEP | abc -> abc |
| R12 | G6P -> F6P | abcdef -> abcdef |
| R13 | DHAP -> GAP | abc -> abc |
| R14 | GAP -> PG3 | abc -> abc |
| R15 | FBP -> DHAP + GAP | abcdef -> abc + def |
| R16 | Pyr.m -> AcCoA.m + CO2 | abc -> ab + c |
| R17 | Pyr.m + CO2 -> OAA.m | abc + d -> abcd |
| R18 | OAA.m + AcCoA.m -> Cit.m | abcd + ef -> abcdef |
| R19 | aKG.m -> Suc.m + CO2 | abcde -> abcd + e |
| R20 | aKG.m -> aKG | abcde -> abcde |
| R21 | Cit.m -> aKG.m + CO2 | abcdef -> abcde + f |
| R22 | Suc.m -> Fum.m | abcd -> abcd |
| R23 | Fum.m -> Mal.m | abcd -> abcd |
| R24 | Mal.m -> OAA.m | abcd -> abcd |
| R25 | Gln.x -> Gln | abcde -> abcde |
| R26 | Gln -> Glu | abcde -> abcde |
| R27 | aKG -> Glu | abcde -> abcde |
| R28 | Asp.m + Mal -> Asp + Mal.m | abcd + efgh -> abcd + efgh |
| R29 | OAA.m + Glu.m -> Asp.m + aKG.m | abcd + efghi -> abcd + efghi |
| R30 | OAA -> Mal | abcd -> abcd |
| R31 | Cit.m -> Cit | abcdef -> abcdef |
| R32 | Cit -> aKG + CO2 | abcdef -> abcde + f |
| R33 | Cit -> AcCoA + OAA | abcdef -> ab + cdef |

**Supplemental Table 4. Network structure for metabolic flux analysis.**

**Statistical Analysis.**

Statistical analysis was conducted using GraphPad Prism software and R studio. Differences between groups were considered significant at $p<0.05$. Sample sizes were not predetermined based on statistical power calculations. Formal randomizations of rat or human heart tissue experiments were not used. Data were tested for normal distribution and similar variance among treatments using the Shapiro-Wilk tests. Statistical significance was calculated by multiple unpaired t-tests and comparisons analysis using a false discovery rate < 5% by the two-step method of Benjamini, Krieger, and Yekutieli, 2-way ANOVA followed by multiple comparison analysis by Sidak, or Mann–Whitney or Wilcoxon tests (when a nonparametric test was appropriate).

**SUPPLEMENTARY FIGURES AND FIGURE LEGENDS**

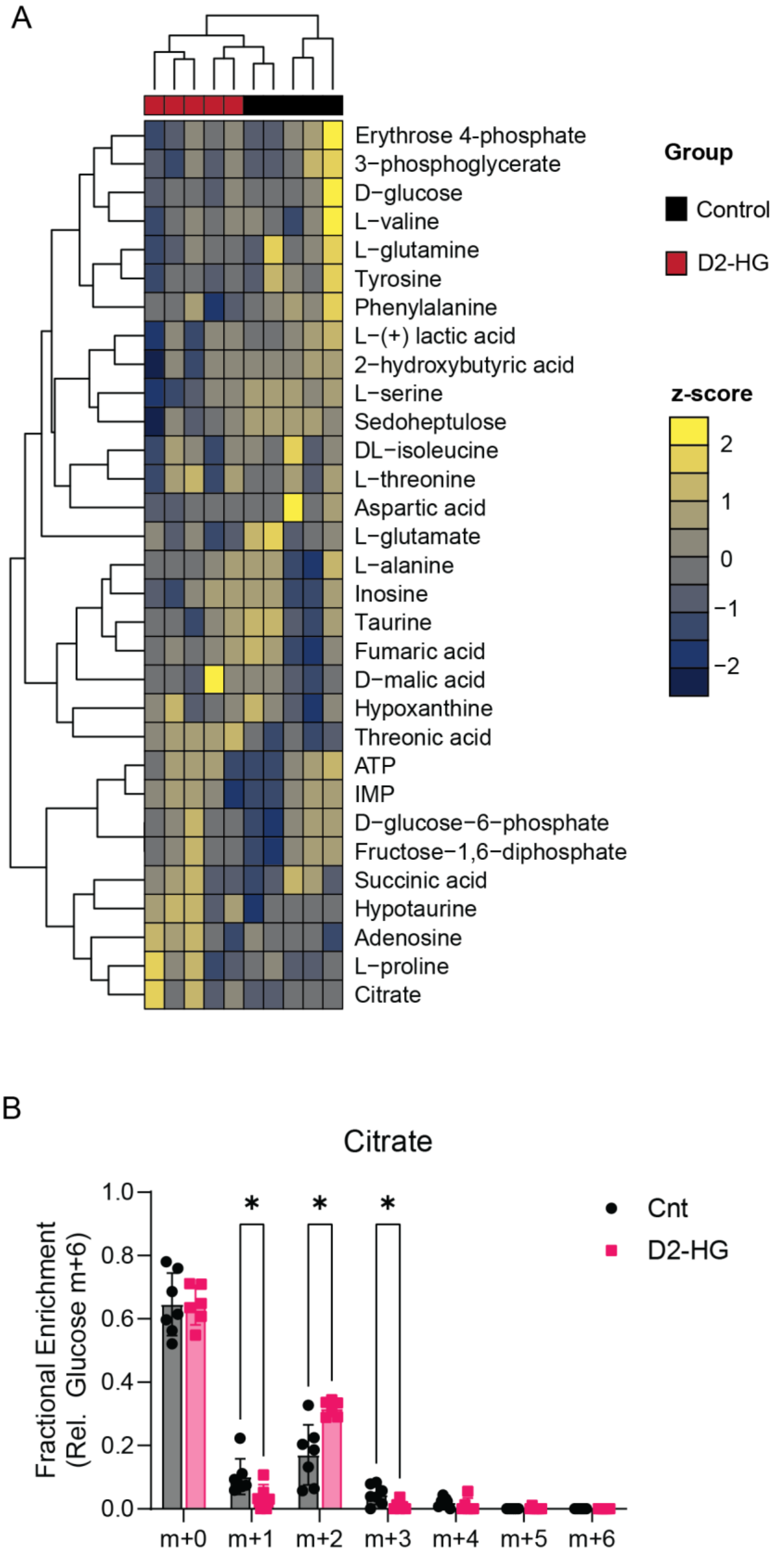


**Supplementary Figure 1. Metabolic characteristics in human heart tissue treated with or without D2-HG.** (**A**) Scaled MS-abundances of metabolites in heart tissue slices from 5 patients treated with or without D2-HG (1.0 mM) for 24 h. Tissue slices from heart donor patients were subjected to D2-HG or PBS (control) allowing the comparison of oncometabolic features within and between patients. (**B**) Isotopologue distribution of citrate in human tissue slices from 6-7 patients treated with or without D2-HG (1.0 mM) and labelled with [U-$^{13}$C]glucose. Statistical analysis was conducted using Welch t-test with FDR<5%. *p-value<0.05.

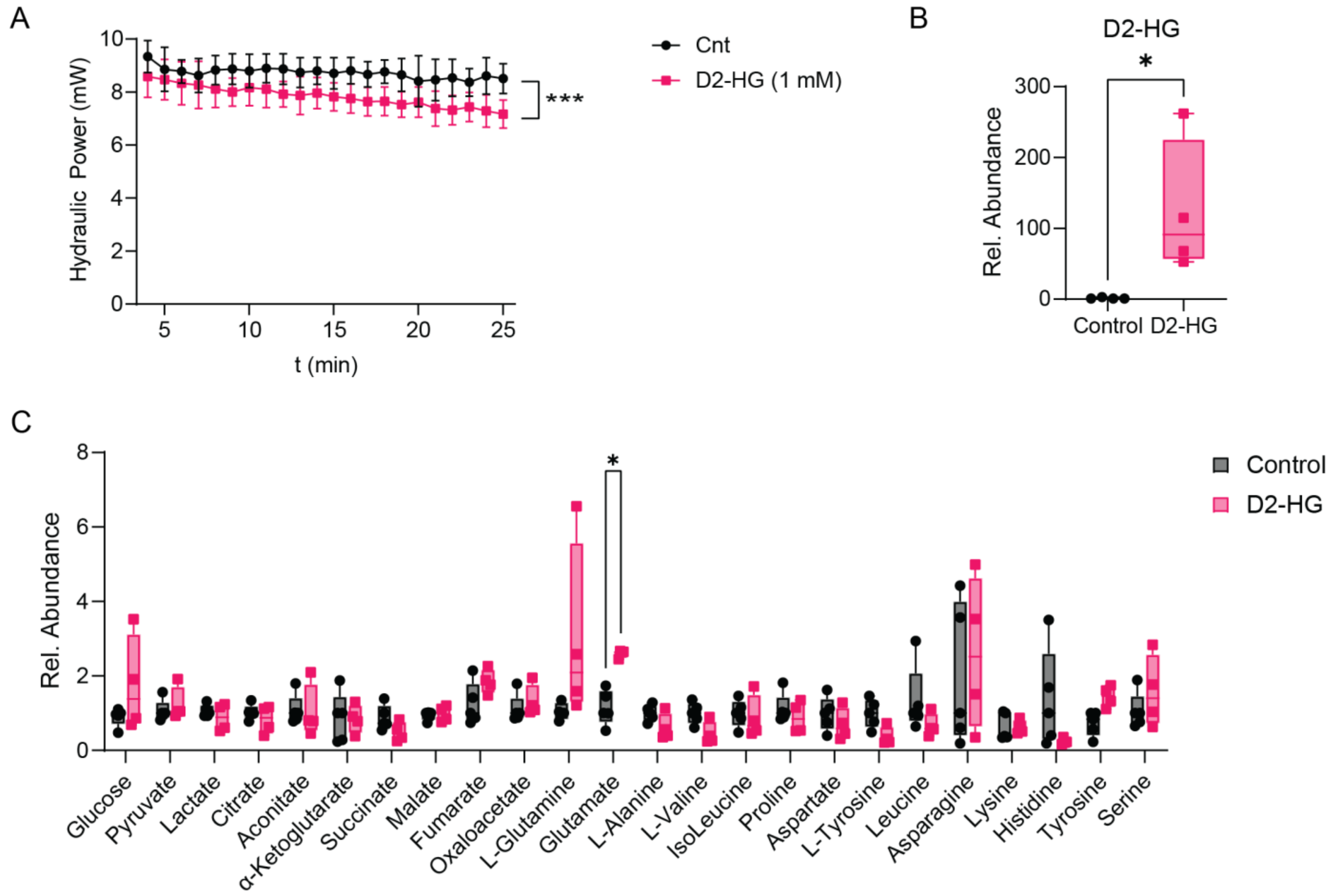


**Supplementary Figure 2. Hydraulic power and metabolic alterations in working rat heart perfusions.** (**A**) Hydraulic power (mW) of rats perfused using the isolated working heart preparations with or without D2-HG (1.0 mM) and physiological concentrations of glucose (5.5 mM), glutamine (0.1 mM), pyruvate (0.5 mM) and oleate (0.1 mM). Statistical analysis was conducted using Kruskal-Wallis test with FDR<5%. N=8-10 rates/group. *p-value<0.05, ***p-value<0.005. (**B**) D2-HG abundance in perfused rat hearts. Statistical analysis was conducted using Welch t-test followed by multiple comparison analysis using Benjamini, Krieger, Yekutieli with FDR<5%. N=4 rates/group. *p-value<0.05. (**C**) Relative abundance of metabolites in rat hearts perfused with or without D2-HG (1.0 mM) for 30 min. Statistical analysis was conducted using Kruskal-Wallis test with FDR<5%. N=4-5 rats/group. *p-value<0.05, ***p-value<0.005.

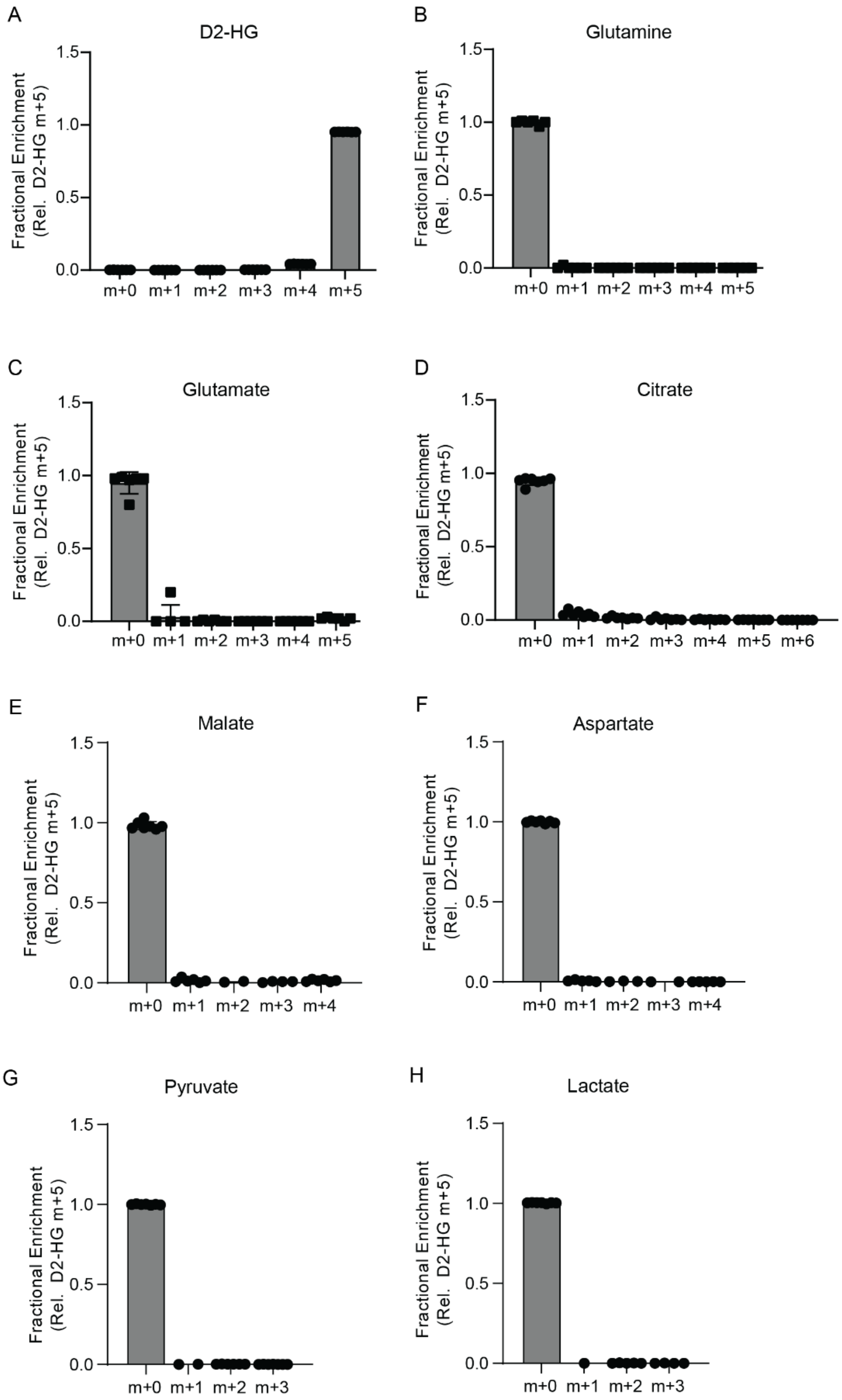


**Supplementary Figure 3. Carbon enrichment in [U-$^{13}$C]D2-HG-perfused rat hearts.** (**A**) Isotopomer distribution of [U-$^{13}$C]D2-HG from rat hearts perfused with D2-HG. (**B-H**) Fractional enrichment of glutamine (**B**), glutamate (**C**), citrate (**D**), malate (**E**), aspartate (**F**), pyruvate (**G**), and lactate (**H**) from rat hearts perfused with [U-$^{13}$C]D2-HG. N=6 rats. Statistical analysis was conducted using Kruskal-Wallis test with FDR<5%. N=4-5 rats/group. *p-value<0.05, ***p-value<0.005.

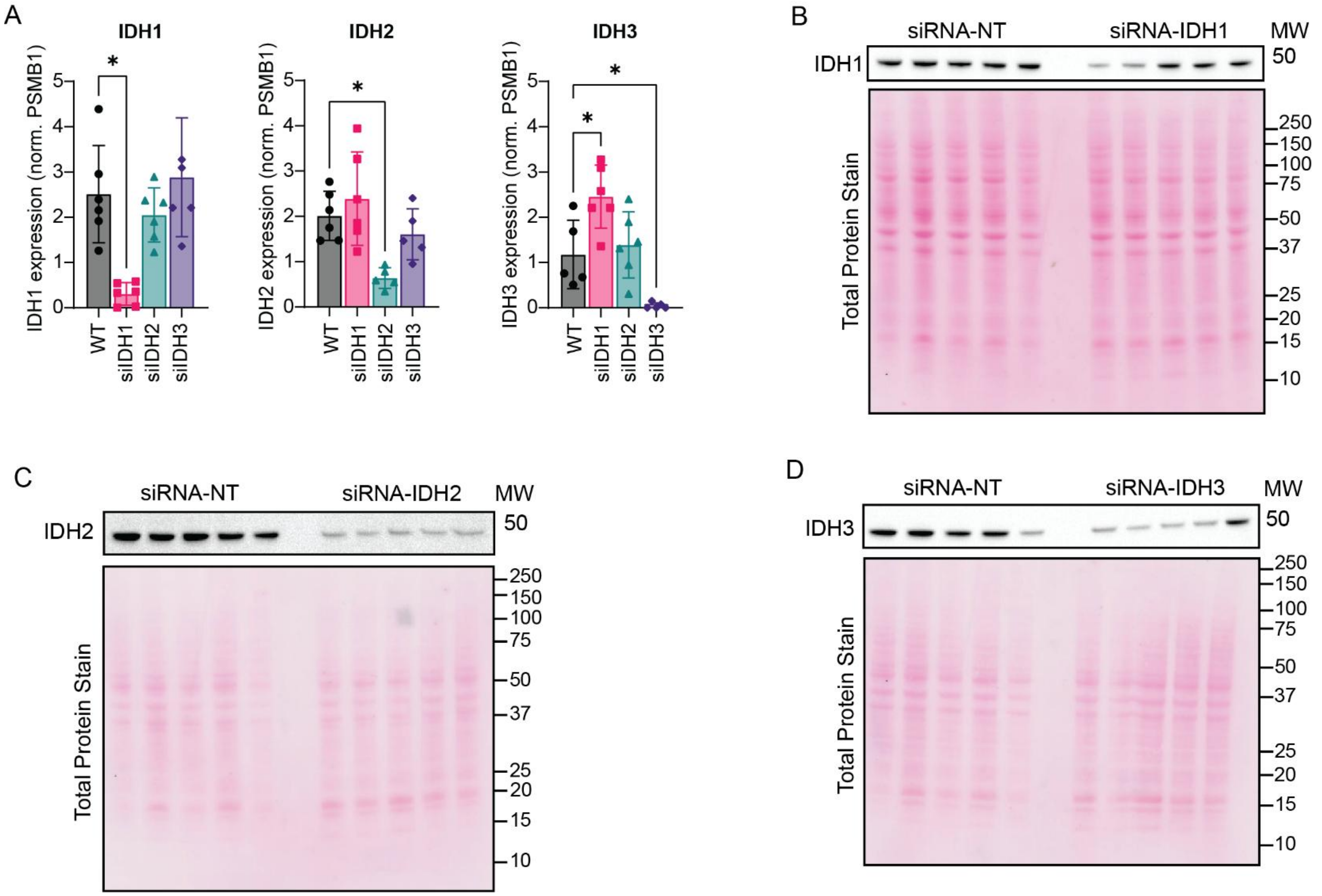


**Supplementary Figure 4. IDH1 isoform drives reductive carboxylation in human cardiomyocytes.** (**A**) Gene expression quantification in differentiated AC16 cells treated with non-targeting silencing RNA (siRNA-NT) and siRNA targeting IDH-isoforms (IDH1, IDH2 and IDH3). Expression was normalized to PSMB1. Quantification was conducted in 5-6 independent experiments. Statistical analysis was conducted using Brown-Forsythe and Welch ANOVA tests and multiple comparisons analysis with Dunnetts T3. *p-value<0.05, **p-value<0.01, ***p-value<0.005, ****p-value<0.001. (**B-D**) Westernblotting of IDH1 (**B**), IDH2 (**B**) and (IDH3) (**D**) in differentiated AC16 cells treated with siRNA-NT or siRNA targeting IDH1, IDH2 and IDH3, respectively. Immunoblots were repeated twice with similar results confirming the successful reduction in IDH-isoform protein expression.
